# OsNAC2 maintains the homeostasis of immune responses to bacterial blight through the OsEREBP1 in rice

**DOI:** 10.1101/2023.08.04.552043

**Authors:** Qun Zhong, Jiangtao Yu, Yiding Wu, Xuefeng Yao, Xiangzong Meng, Feng Ming

**Author notes:** These authors contributed equally to this work (Qun Zhong, Jiangtao Yu). Corresponding author: Feng Ming.

## Abstract

Rice bacterial blight, caused by *Xanthomonas oryzae* pv. *Oryzae* (*Xoo*), threatens plant growth and yield. However, the molecular mechanisms underlying rice immunity against *Xoo* remain elusive. Here, we demonstrated down-regulation of a NAC (NAM-ATAF-CUC) transcription factor *OsNAC2* enhanced resistance to bacterial blight disease in rice. Consistently, salicylic acid (SA) biosynthesis was greatly attenuated when OsNAC2 overexpressed. Furthermore, study demonstrated that OsNAC2 can down-regulated the expression of SA synthesis genes, thus mediating SA signal transmission. Simultaneously, OsNAC2 interacted with OsEREBP1 (AP2/ERF family) in the nucleus, may be bloking the inhibition of OsNAC2 to *OsICS1*, *OsPAL3* and so on. Furthermore, OsEREBP1 interacted with OsXb22a in the cytoplasm to exert its positive regulatory effects to bacterial blight. However, *OsNAC2*-overexpression kept OsEREBP1 in the nucleus and be rapidly degraded in pathogen infection, which adversely affects the interaction of OsEREBP1-OsXb22a. Our results determined that OsNAC2 inhibits the SA signaling and stably interacts with OsEREBP1 to maintain disease resistance. This OsNAC2-OsEREBP1-based homeostatic mechanism provides new insights into rice disease resistance, and it may be useful for improving the disease resistance of important crops through breeding.

## Introduction

Plant diseases profoundly affect global food security and sustainable agricultural development (Jones and Naidu, 2019). Rice (*Oryza sativa*), which is one of the most important food crops, is the potential host of various pathogenic bacteria in the agro-ecosystem. For example, *Xanthomonas oryzae* pv. *oryzae* (*Xoo*), which is a major bacterial pathogen with a high mutation rate, causes a rice disease (bacterial blight) that can spread quickly (i.e., wide incidence), leading to substantial yield losses (Djedatin et al., 2016). Exploiting natural disease resistance genes to breed new rice varieties resistant to bacterial blight is an economical and environmentally friendly strategy for enhancing rice production. The cloning of major resistance genes is an objective in the area of the bacterial blight disease research (Hou et al., 2015). At least 40 bacterial blight resistance genes have been identified, some of which have been cloned and widely used for disease resistance breeding (Oliva et al., 2019; Wang et al., 2017).

In plants, the perception of a signal from an invading pathogen leads to the activation of specific transcription factors via a series of signaling events. The main disease resistance-related transcription factors are bZIP-type TGA family members (Zander et al., 2010), ERF transcription factors (Liu et al., 2012), WRKY family members containing zinc finger domains (Tao et al., 2009; Yang et al., 2009), and NAC family members (Nakashima et al., 2007; Jeong et al., 2010). There is increasing evidence that some NAC transcription factors influence plant immunity through hormone signaling pathways. For example, salicylic acid (SA) is crucial for plant defenses against pathogens (Nawrath and Métraux, 1999) because it mediates the establishment of basic defense responses (Meng et al., 2020), the amplification of immune responses in locally infected tissues (Zavaliev et al., 2020), and the development of systemic acquired resistance (Lawton et al., 1995). Salicylic acid biosynthesis mainly occurs in the chloroplast, with isochorismate synthase (ICS) catalyzing one of the key reactions. Additionally, a small amount of SA is derived from the cinnamic acid produced by phenylalanine ammonia lyase (PAL) in the cytoplasm (M é traux, 2002). In Arabidopsis (*Arabidopsis thaliana*), the NAC transcription factor NTM1-LIKE9 (NTL9) rapidly activates the expression of the SA biosynthesis gene *ICS1* during flg22-triggered immunity (Zheng et al., 2015). The expression of the *CROWDED NUCLEI* (*CRWN*) family *CRWN1* gene is regulated by pathogenic bacteria and SA at the transcriptional and post-transcriptional levels; the encoded protein interacts with NTL9 to enhance the transcriptional inhibition of the downstream disease resistance gene *PR1* (Guo et al., 2017).

In rice, OsEREBP1, which belongs to the AP2/ERF family (163 members), comprises one or two highly conserved AP2 domains and mediates important responses to environmental stimuli or plant growth and development (Sharoni et al., 2011). It also contributes to hormone signal transduction and abiotic stress responses (Chen et al., 2012; Rahimi et al., 2016). A previous study revealed that OsEREBP1 is a signal transduction factor for the specific interaction between the bacterial blight pathogen and rice; lines overexpressing *OsEREBP1* exhibit increased bacterial blight resistance (Jisha et al., 2015). OsEREBP1 can interact with OsXb22a, implying that OsEREBP1 affects rice resistance to bacterial blight (Park et al., 2008; Seo et al., 2011).

Previous research proved that *OsNAC2* helps regulates many biological processes in rice, including root growth and development (Mao et al., 2019), leaf senescence (Mao et al., 2017), and abiotic stress tolerance (Shen et al., 2017). In this study, we determined that *OsNAC2* negatively regulates rice bacterial blight resistance through the SA signaling pathway. Moreover, OsNAC2 interacts with OsEREBP1 in the nucleus, destroying the signination of SA-mediated signialing. In the cytoplasm, OsEREBP1 interacts with OsXb22a to exert its positive regulatory effects. Thus, OsNAC2 balances the distribution of OsEREBP1 in the nucleus and cytoplasm, but it also decreases the OsEREBP1 level during pathogen infections, thereby disrupting immune regulatory homeostasis. The findings of this study provide insights into the maintenance of immune homeostasis.

## Results

### *OsNAC2* plays negative roles in defense response against *Xoo*

We previously demonstrated that *OsNAC2* regulates programmed cell death (Mao et al., 2018), which can protect plants from pathogenic bacteria (Sabater and Martín, 2013). We also observed that the senescence degree of RNAi line was slower than that of Nip at the later stage of grain maturation (Supplemental Fig. S1). To determine whether *OsNAC2* had the role in plant immunity, we examined *OsNAC2* expression in rice leaves inoculated with *Xoo* strain PXO71. Real-time quantitative PCR (RT-qPCR) revealed *OsNAC2* expression was significantly induced by the PXO71 infection compared with the control group, with an approximately 4.5-fold increase at 2 h post-inoculation (Fig. 1A). We infected the previously constructed *OsNAC2*-transgenic lines and observed the phenotype by *Xoo* strain PXO71 (Chen et al., 2015). Compared with the wild-type (WT), the *OsNAC2*-overexpressing lines (ON7 and ON11) were substantially less resistant to PXO71, but the *OsNAC2*-RNAi lines (RNAi25 and RNAi31) were significantly more resistant (Fig. 1, B and C). NBT and DAB staining analysis that the RNAi25 and RNAi31 leaves produced more H_2_O_2_ and O_2_^-^ than the ON7 and ON11 after infection (Fig. 1D). In addition, the expression trend of resistance marker genes *OsPR1a* and *OsPR10a* was consistent with the disease resistance phenotype (Fig. 1, E and F). These results indicated that *OsNAC2* negatively regulates bacterial blight in rice.

**Figure 1.**
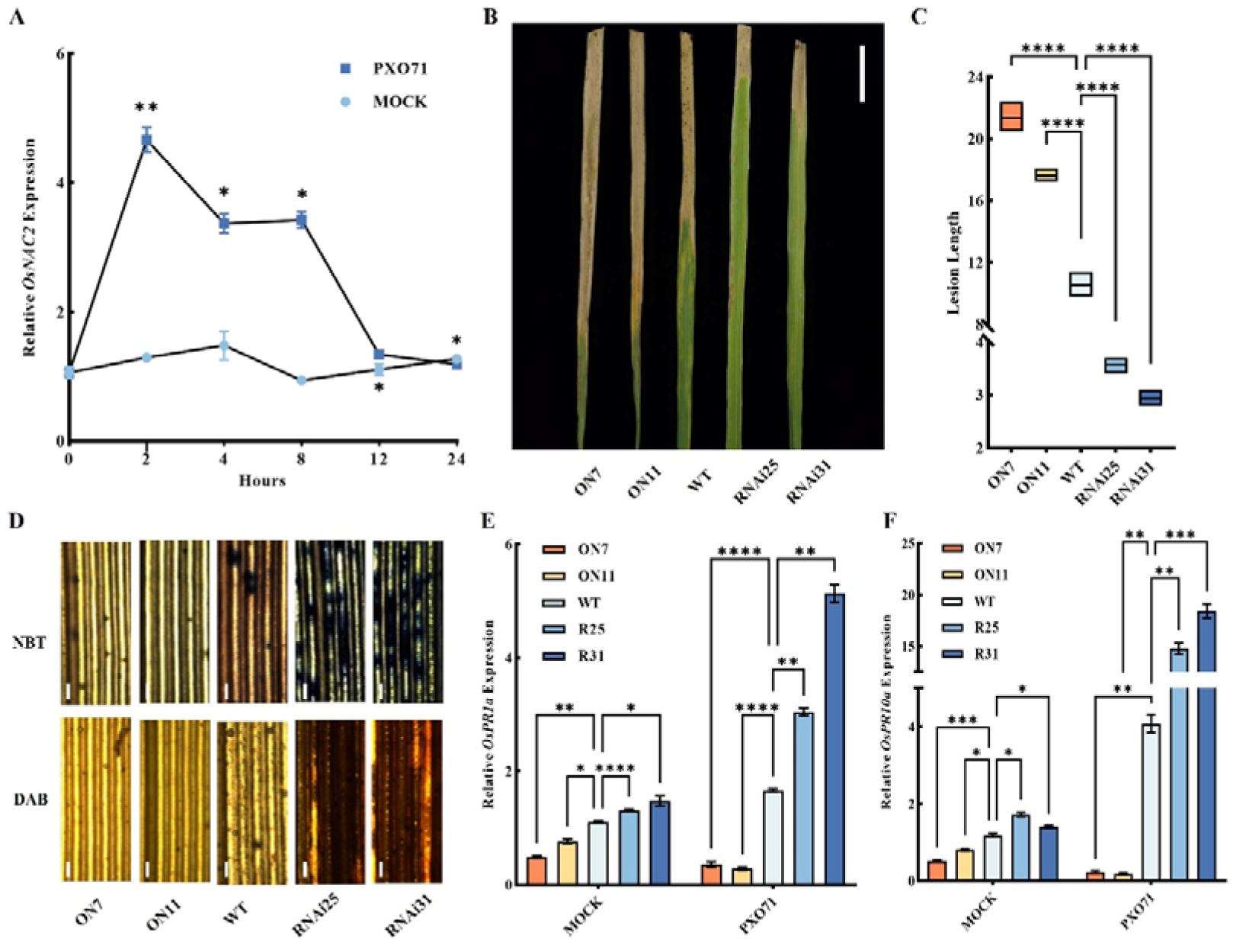
O*s*NAC2 negatively regulates rice immunity against *Xoo* infection. A, Expressions of *OsNAC2* gene in the WT seedlings under the *Xoo* strain PXO71 infection at different time periods. B-C, Disease phenotype (B) and lesion length (C) of the WT and *OsNAC2*-transgenic lines after 4 weeks of infect with PXO71 under natural conditions. Scale bar=5 cm D, NBT or DAB staining of the WT and *OsNAC2*-transgenic lines leaves after 4 weeks of infect with PXO71 under natural conditions. Scale bars=100 µm. E-F, Expressions of *OsPR1a* (E) and *OsPR10a* (F) genes in WT and *OsNAC2*-transgenic lines under the PXO71 infection at different time periods. Data are presented as the mean ± standard error of at least three biological replicates. Asterisks indicate significant differences between treatment and control by *t*-test. *P< 0.05, **P< 0.01, and ***P< 0.001.

### OsNAC2 inhibites the accumulation of SA in vivo

AtNPR1 is a transcriptional activator, and as a receptor for SA, the binding of SA increases its transcriptional activation activity, leading to enhanced plant resistance (Ding et al., 2018). Furthermore, the NAC transcription factor family may be involved in the transcriptional activation of genes in the SA signaling pathway in *Arabidopsis* (Zheng et al., 2012). Thus, we wondered whether *OsNAC2* is involved in the SA pathway to regulate plant immunity. To validate this hypothesis, we first analyzed the *OsNAC2* expression in the WT with or without treatment with SA. Compared with the Mock, *OsNAC2* expression was significantly increased by approximately 3-10-fold in the WT treatment with exogenous 100 µM SA (Fig. 2A). Moreover, the SA contents of the RNAi31 was 30.08 nl/g.h, which was significantly higher than that of the WT (25.02 nl/g.h), while the ON7 showed the lowest SA emission (27.83 nl/g.h) (Fig. 2B).

**Figure 2.**
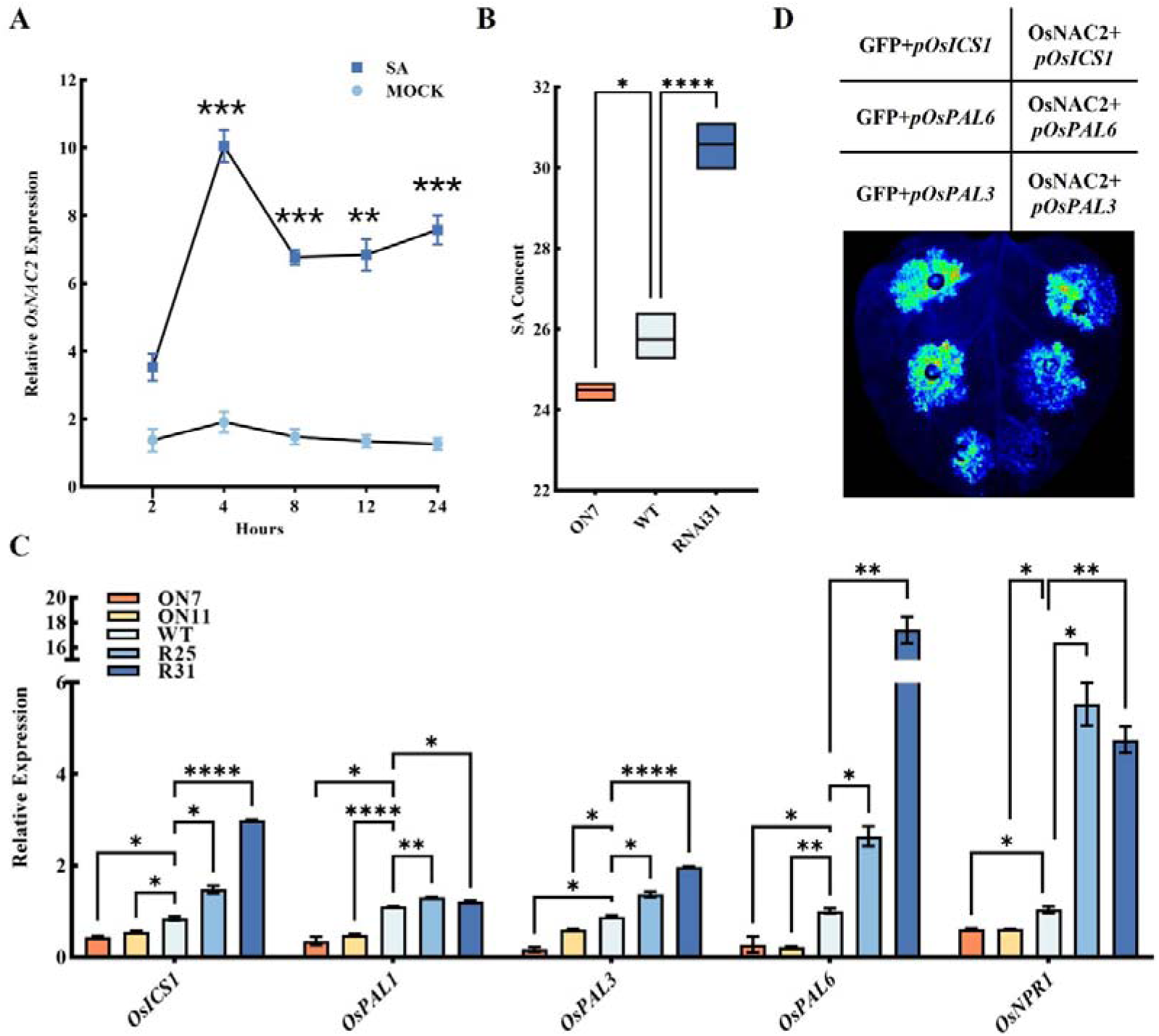
O*s*NAC2 regulates the salicylic acid signaling pathway. A, Expressions of *OsNAC2* genes in the WT treated with 100 µ mol/L salicylic acid at different time periods. Clean water treatment as the control group. B, SA production in WT, *OsNAC2*-overexpressing line (ON7) and *OsNAC2*-RNAi line (RNAi31). C, Expressions of *OsNPR1* gene in the WT and *OsNAC2*-transgenic lines. C, Expressions of SA biosynthetic genes *OsICS1*, *OsPAL1*, *OsPAL3*, *OsPAL6* and SA receptor *OsNPR1* in the WT and *OsNAC2*-transgenic lines. Error bars represent the standard error of three biological replicates. *p < 0.05, **p < 0.01, and *p < 0.001. D, OsNAC2 inhibits promoter activity of *OsICS1*, *OsPAL3* and *OsPAL6* in *N. benthamiana* leaves. *OsICS1*, *OsPAL3* and *OsPAL6* promoters were fused with LUC reporter. *35S:* empty vector served as a negative control.

To further identify whether *OsNAC2* directly regulated genes involved in the SA metabolism or signaling, we examined the expression levels of SA synthesis genes *OsICS1* and *OsPALs* genes. The *OsICS1*, *OsPAL3* and *OsPAL6* genes expression were sharply increased in the RNAi25 and RNAi31 lines than in the WT, while were distinctly decreased in the ON7 and ON11, including *OsPAL1* (Fig. 2C). Subsequently, we verified whether OsNAC2 binds to the promoter of those genes. As shown in Figure 2D, we demonstrated that OsNAC2 inhibited the transcription of *OsICS1*, *OsPAL3*, and *OsPAL6* using the double transfer activated luciferase system (LUC/REN) in *N. benthamiana* leaves. When co-infiltration of *35S:OsNAC2* and promoter-*OsICS1*, constructs led to a weaker luciferase reporter signal, while *35S:* empty vector does not. The same is true for *OsPAL3* and *OsPAL6* promoters. Therefore, OsNAC2 may indirectly inhibit SA synthesis and regulate plant immunity.

### OsEREBP1 physically associates with OsNAC2 in the nucleus to inhibit the transcription of OsNAC2

To further understand how OsNAC2 affects plant immunity and identify the potential proteins of OsNAC2, we extracted proteins from the ON11 were immunoprecipitated using an anti-GFP antibody. The bands corresponding to the differentially abundant proteins between the ON7 and WT lines were excised from gels for a mass spectrometry analysis. There were 927 differentially abundant proteins in the *OsNAC2*-overexpressing lines (ON11) compared to the WT (Supplemental Fig. S2A). Because *OsNAC2* negatively regulates rice resistance to bacterial blight, according to the correlation degree and function of proteins, we finally found 7 candidate genes (Supplemental Fig. S2B), and only 3 genes *OsSGT1* (LOC_Os01g0624500), *OsDSG1* (LOC_Os06g0154500), and *OsEREBP1* (LOC_Os02g54160) may be involved in plant immunity-related processes (Supplemental Fig. S2C). Further study proved that the expression of *OsEREBP1* was consistent with the resistance phenotype of OsNAC2 in *OsNAC2*-transgenic lines (Supplemental Fig. S3), so *OsEREBP1* may interacts with OsNAC2.

In Y2H assays, OsNAC2 interacted strongly with OsEREBP1, whereas the corresponding control cells were unable to grow on this medium (Fig. 3A). We then verified the OsNAC2 co-immunoprecipitated with OsEREBP1 only when OsNAC2-GFP-HA and OsEREBP1-GFP-Myc were co-expressed in *N. benthamiana* leaves (Fig. 3B). In addition, bimolecular fluorescence complementation (BiFC) assay showed that the fluorescence signal was observed in the nucleus of *N. benthamiana* leaves co-infiltrated with *OsNAC2*-nYFP and *OsEREBP1*-cYFP, which co-localized with the nucleus marker mCherry (Fig. 3C). These results suggest that OsNAC2 and OsEREBP1 may play an important role in the nucleus. We therefore selected OsEREBP1 for further exploration.

**Figure 3.**
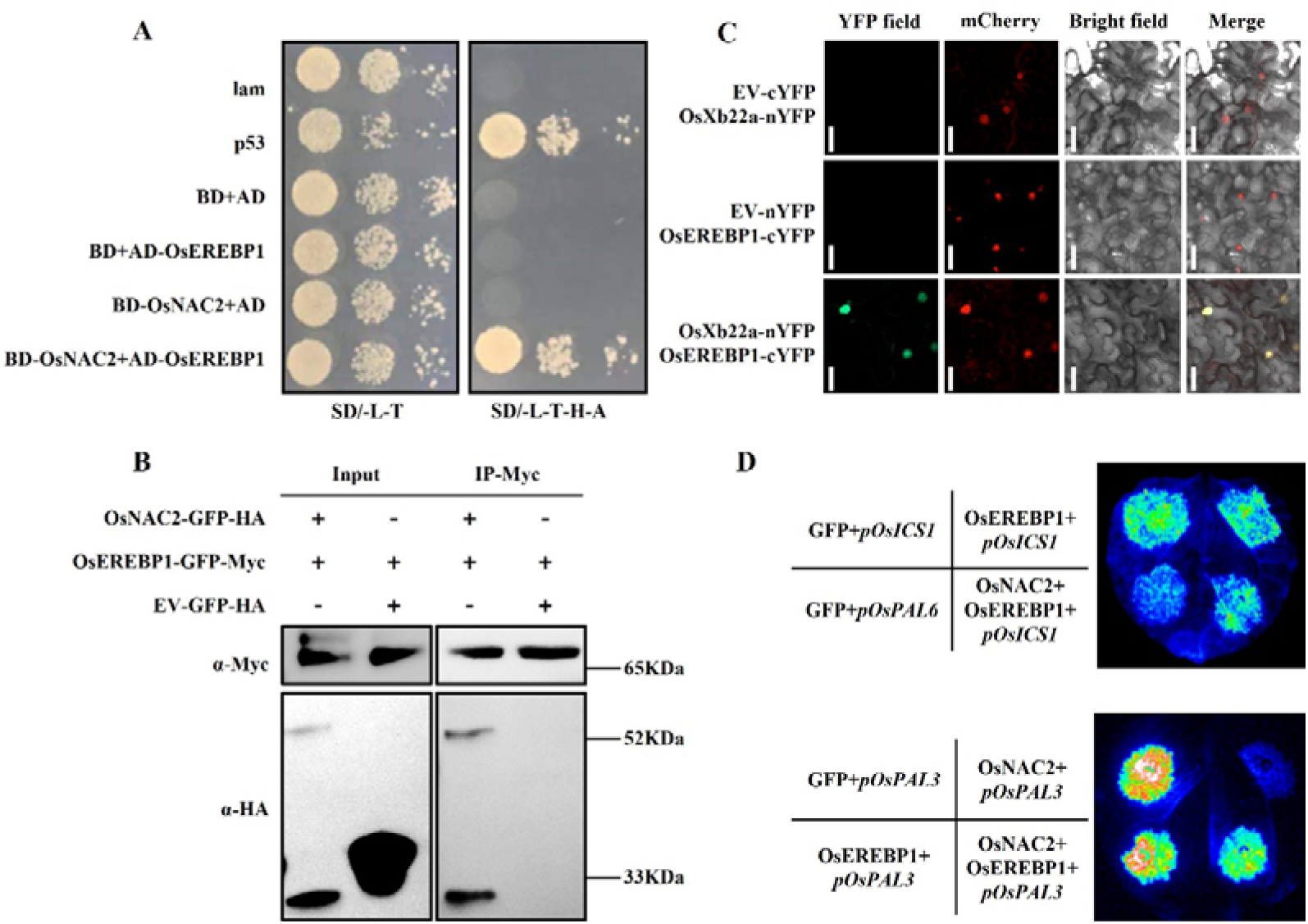
OsNAC2 interacts with OsEREBP1 protein in the nucleus. A, Y2H assays of the interaction between OsNAC2 and OsEREBP1. Transformed yeast cells were diluted in three gradients and grown on SD/-T/-L and SD/-T/-L/-H/-A medium. B, Co-immunoprecipitation (Co-IP) analysis of the interaction between OsNAC2 and OsEREBP1 in *N. benthamiana* leaves transformed with *35S*: OsNAC2-GFP-HA with *35S*: OsEREBP1-GFP-Myc. *35S*: GFP-HA served as the control. The experiment was repeated three times with similar results. C, Bimolecular fluorescence complementation (BiFC) analysis showed the interaction between OsNAC2 and OsEREBP1 in *N. benthamiana* leaves. The plasmids carrying the structures of *OsNAC2*-*n*YFP and *OsEREBP1*-*c*YFP together were transiently expressed in *N. benthamiana* leaves. At 48h after transformation, fluorescence was observed using a confocal microscope. The experiment was repeated three times with similar results. Scale bars=500 µm. D, OsEREBP1 undermines OsNAC2’s suppression on the *OsICS1* or *OsPAL3*-targeted gene expression. Transient luciferase activities in *N*. *benthamiana* were analyzed by co-transforming with the LUC reporter and different combinations of effectors. LUC gene is driven by the *OsICS1* and *OsPAL3* promoter. GFP serves as the negative control.

### OsEREBP1 undermines OsNAC2’s suppression on the SA synthetic genes expression

These results suggested that OsNAC2 and OsEREBP1 might form a complex to co-regulate the expression of target genes. To test this idea, we selected the *OsICS1* and *OsPAL3* genes as a gene potentially regulated by OsNAC2 and OsEREBP1, as the *ICS1* promoter contains TAGC that is putative binding sites for NAC, while there is no ERF-bound GCCGCC element, *PAL3* promoter contains both TAGC and GCCGCC (Supplemental Fig. S4). So, we investigate whether OsEREBP1 could affect OsNAC2-mediated genes transcription, using the firefly luciferase (LUC) reporter gene driven by the *ICS1* and *PAL3* promoters and OsNAC2 and OsEREBP1 as effectors, we observed that OsNAC2 alone repressed transcription from the *OsICS1* and *OsPAL3* (Fig. 2D), whereas those transcriptional reductions were recovered to varying degrees when *OsNAC2* and *OsEREBP1* were co-expressed (Fig. 3D). These results indicated that OsNAC2 and OsEREBP1 act synergistically to regulate SA signaling.

### OsEREBP1 stabilizes OsXb22a by interacting with it in the cytoplasm

As a positive regulator of rice responses to bacterial pathogens, OsEREBP1 interacts with the disease resistance-related protein OsXb22a (Jisha et al., 2015). Further studies showed that, through Y2H and Co-IP assays, yeast cells containing BD-OsXb22a and AD-OsEREBP1 grew well on the SD/-T-L-H-A, whereas the corresponding control cells were unable to grow on this medium (Fig. 4A). Furthermore, OsXb22a coimmunoprecipitated with OsEREBP1 only when OsXb22a-GFP-Flag and OsEREBP1-GFP-Myc were co-expressed in *N. benthamiana* leaves (Fig. 4B). These results suggested that OsXb22a interacts with OsEREBP1. Interestingly, for bimolecular fluorescence complementation (BiFC) assay, we found that the fluorescence signal presented in the cytoplasm of *N. benthamiana* leaves co-infiltrated with OsXb22a-nYFP and OsEREBP1-cYFP, the YFP signial nonlocated with *m*Cherry (Fig. 4C). Thus, OsXb22a was co-localized with OsEREBP1 in the cytoplasm. Notably, this is unlike the OsNAC2 interacts with OsEREBP1 in the nucleus (Fig. 3, A-C). In addition, through western blot analysis indicated that the co-expression of OsEREBP1 and OsXb22a resulted in the increase of OsXb22a protein level, while the presence of the protease inhibitor MG132 made this increase even more obvious. (Fig. 4D). Together, these results indicated that OsEREBP1 interacts with OsXb22a in the cytoplasm and stabilizes OsXb22a.

**Figure 4.**
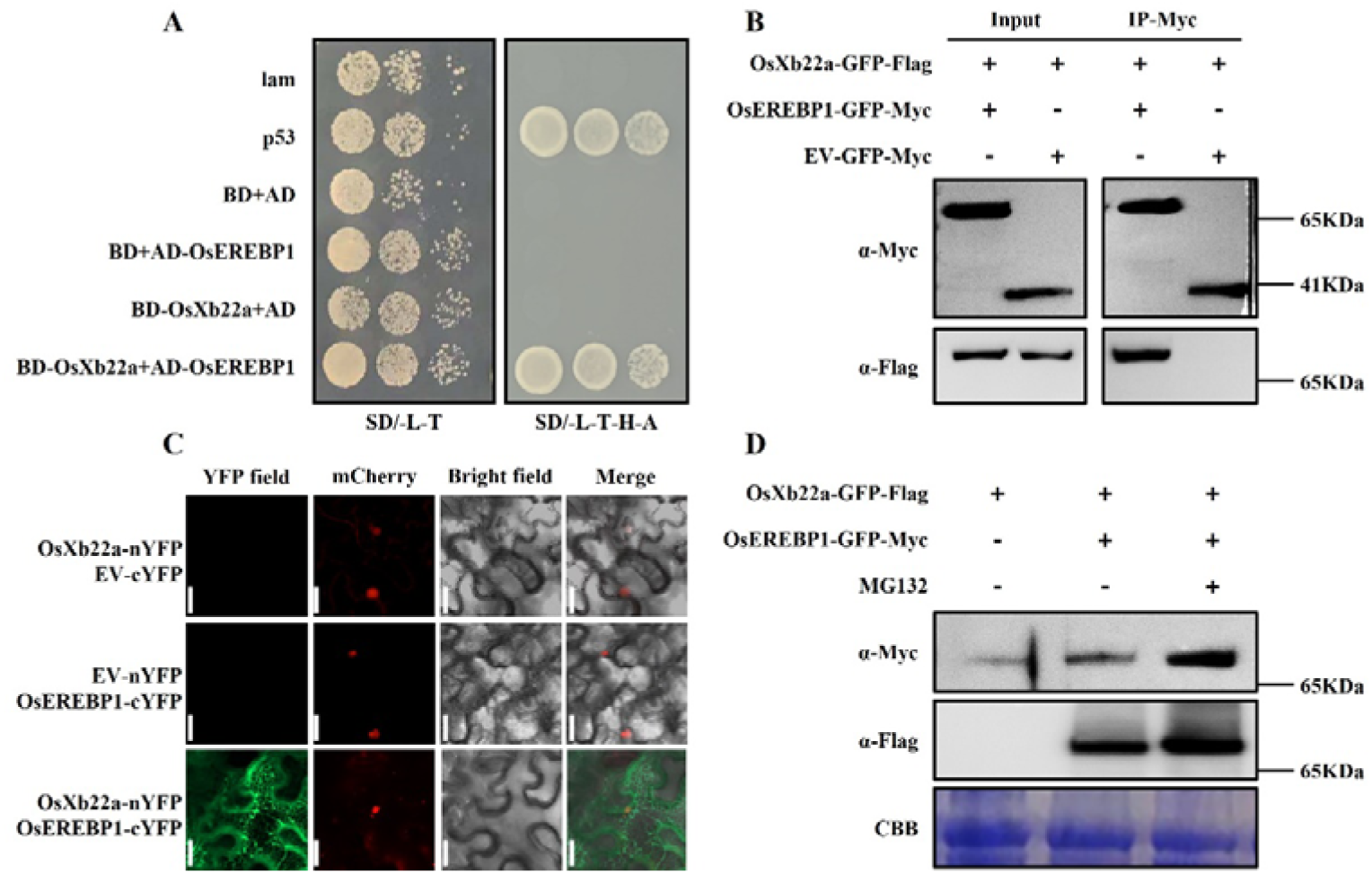
OsEREBP1 interacts with OsXb22a in the cytoplasm. A, Yeast two-hybrid analysis of the interaction between OsXb22a and OsEREBP1. Transformed yeast cells were diluted in three gradients and grown on SD/-T-L and SD/-T-L-H-A medium. B, Co-immunoprecipitation (Co-IP) analysis of the interaction between OsXb22a and OsEREBP1 in *N. benthamiana* leaves transformed with 35S-OsXb22a-GFP-Flag with 35S-OsEREBP1-GFP-Myc. 35S-GFP-Myc served as the control. The experiment was repeated three times with similar results. C, Bimolecular fluorescence complementation (BiFC) analysis showed the interaction between OsXb22a and OsEREBP1 in *N. benthamiana* leaves. The plasmids carrying the structures of *OsXb22a*-nYFP and *OsEREBP1*-cYFP together were transiently expressed in *N. benthamiana* leaves. At 48h after transformation, fluorescence was observed using a confocal microscope. The experiment was repeated three times with similar results. Scale bars=500 µm. D, Western Blot analysis of the interaction between OsXb22a and OsEREBP1 in *N. benthamiana* leaves transformed with 35S-OsXb22a-GFP-Flag with 35S-OsEREBP1-GFP-Myc and protease inhibitor (MG132). OsXb22a-GFP-Flag served as the control. The bottom panel shows staining with Coomassie Brilliant Blue (CBB) as a loading control.

### OsNAC2 imprisons OsEREBP1 in the nucleus

We considered it was necessary to determine the subcellular localization of OsNAC2, OsEREBP1 and OsXb22a owing to the fact that the cellular compartment of OsEREBP1 interacts with OsNAC2 or Osxb22a were not identical. Therefore, we performed the subcellular localization of OsNAC2, OsXb22a and OsEREBP1 in rice protoplasts. This result showed that OsNAC2 was localized in the nucleus and OsXb22a was localized in the cytoplasm, whereas OsEREBP1 was located in these two places (Fig. 5A). Interestingly, when OsEREBP1-YFP and OsNAC2 were co-expressed, most of the fluorescence signal of OsEREBP1 was concentrated in the nucleus in rice protoplasts, co-located with the nuclear marker *mCherry*. However, when OsEREBP1-YFP and OsXb22a were co-expressed, they were concentrated in the cytoplasm in rice protoplasts (Fig. 5A). The results were also verified in *N. benthamiana* leaves (Fig. 5B), namely, OsNAC2 enabled OsEREBP1 to be located in the nucleus, and OsXb22a enabled OsEREBP1 to be transferred to the plasmid. These results indicated that OsNAC2 or OsXb22a clearly affected the localization of OsEREBP1, and the interaction between OsNAC2 and OsEREBP1 inhibited its exit from the nucleus.

**Figure 5.**
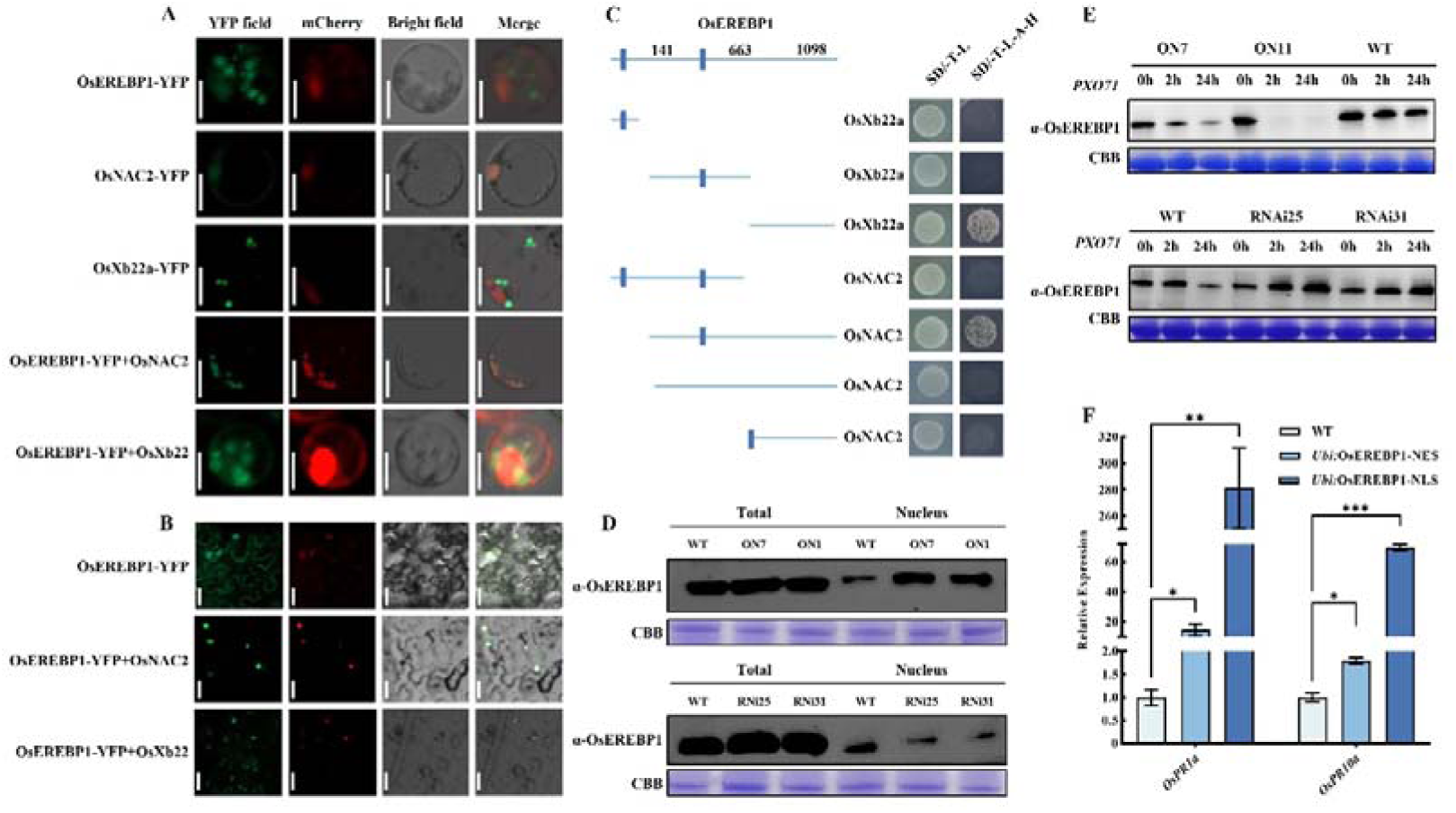
OsNAC2 restricts OsEREBP1 in the nucleus and degrades it during the process of pathogen infection. A, OsNAC2 induces OsEREBP1 to locate in the nucleus. OsNAC2, OsXb22a and OsEREBP1-YFP were transiently expressed in rice protoplasts. *mCheery* was used as nuclear marker. Confocal microscope was used to observe the OsEREBP1-YFP subcellular localization. Bar=500 µm. B, Subcellular localization of OsEREBP1 is altered by overexpression of OsNAC2 or OsXb22a in *N. benthamiana* leaves transformed with *35S:*OsEREBP1-YFP with *35S:*OsNAC2, *35S:*OsEREBP1-YFP with *35S:*OsXb22a. *35S:*OsEREBP1-YFP served as the control. The experiment was repeated three times with similar results. Bars = 100 µm. C, Yeast two-hybrid assays of the interaction between OsEREBP1 and OsNAC2 or OsXb22a. Truncated based on the nuclear localization signal domain found in OsEREBP1, the specific interaction regions with OsNAC2 and OsXb22a were determined by yeast two-hybrid experiment. Transformed yeast cells were diluted in three gradients and grown on SD/-T/-L and SD/-T/-L/-H/-A medium. D, Western blot analysis of OsEREBP1 protein levels in nuclear extracts from *OsNAC2*-transgenic lines. Total protein and nuclear protein were extracted from two-week-old rice seedlings, the nuclear protein was separated by the method of *Nuclear and Cytoplasmic Extract Protocol* (Baldwin, 1996). The protein was detected by immunoblotting with α-OsEREBP1 antibody. This experiment was repeated three with similar results. The bottom panel shows staining with Coomassie Brilliant Blue (CBB) as a loading control. E, Western blot analysis of OsEREBP1 protein levels in the WT and *OsNAC2*-transgenic lines under the *Xoo* strain PXO71 infection at different time periods. The protein was detected by western blotting with α-OsEREBP1 antibody. Uninoculated bacterial blight served as the control. The bottom panel shows staining with Coomassie Brilliant Blue (CBB) as a loading control. F, Detection of *Ubi*: OsEREBP1-NLS and *Ubi*: OsEREBP1-NES resistance gene expression after inoculation. Asterisks indicate significant differences between treatment and control by *t*-test. *P< 0.05, **P< 0.01, and ***P< 0.001.

To investigate the specific sites where OsNAC2 and OsXb22a interact with OsEREBP1, we analyzed the conserved domain of OsEREBP1 and truncated the OsEREBP1 fragment according to the conserved domain. Through yeast two-hybrid assays, we observed that OsNAC2 mainly interacts with the 142-1098 region of OsEREBP1, which contains an N-terminal nuclear localization sequence and the full-length C-terminal, whereas OsXb22a interacts with the C-terminal domain (664-1098) that lacking a nuclear localization sequence. To assess whether the second nuclear localization sequence of *OsEREBP1* determines the interaction between OsNAC2 and OsEREBP1, we fused a nuclear export signal (NES) sequence to the N-terminal end of OsEREBP1^142-1098^ (OsEREBP1^142-1098^NES). Moreover, a nuclear localization sequence was fused to the N-terminal end of OsEREBP1^664-1098^ (OsEREBP1^664-1098^NLS). Unfortunately, with the addition of NLS and NES, OsEREBP1 did not interact with OsNAC2. These results indicated that OsNAC2 and OsXb22a interact with OsEREBP1 in different segments, and OsEREBP1second localization signal is apparently important for the interaction with OsNAC2 (Fig. 5C). Next, we confirmed the localization of OsNAC2-OsEREBP1 in the nucleus by examining nuclei isolated from seedling cells using an antibody that specifically recognizes OsEREBP1. This analysis provided further evidence of the localization of OsEREBP1 in the nucleus and cytoplasm of WT cells (Supplemental Fig. S5). However, there was more OsEREBP1 in the nuclei of ON7 and ON11 plants than in the nuclei of RNAi25 and RNAi31 plants (Fig. 5D), suggesting that the increased expression of OsNAC2 promoted the retention of OsEREBP1 in the nucleus.

### OsNAC2 can degrade OsEREBP1 when infected with pathogen

We explored whether this change in protein distribution of OsEREBP1 affects the disease resistance mechanism. At 2 h and 24 h post-inoculation with PXO71, OsEREBP1 was almost completely degraded in the ON7 and ON11 plants, while degraded less in WT. In contrast, OsEREBP1 accumulated significantly at 2 h and 24 h post-inoculation in the RNAi25 and RNAi31 lines (Fig. 5E). These findings further illustrated the roles of OsEREBP1 and OsNAC2 in rice responses to bacterial blight. More specifically, PXO71 induced the expression of OsNAC2, leaving OsEREBP1 in the nucleus and then degrading, thereby limiting the positive regulatory function of OsEREBP1. If it is true that OsNAC2 degrades into nuclear OsEREBP1 when infected by pathogenic bacteria, as we assumed, the stable expression of OsEREBP1 in the nucleus will greatly reduce disease resistance. In order to verify this, we generated *Ubi*: OsEREBP1-NLS and *Ubi*: OsEREBP-NES transgenic lines. identified 14 transgenic positive seedlings of *Ubi*: OsEREBP1-NLS and 13 transgenic positive seedlings of *Ubi*: OsEREBP1-NES (Supplementary Fig. S6A), and four of them were selected for resistance detection with no significant difference in OsEREBP1 expression (Supplemental Fig. S6B). The results showed that the expression of *PRs* were significantly induced in OsEREBP-NLS after inoculation, but the degree of induction in OsEREBP-NES was less than that of OsEREBP-NLS lines (Fig. 5F). These results indicate that OsEREBP1 actively functions in the nucleus as a positive immune regulator.

### The resistance of *osnac2 oserebp1* mutant lines decreased

To elucidate the genetic relationship between *OsNAC2* and *OsEREBP1* and further functionally characterize OsNAC2 and OsEREBP1 regarding their regulatory effects on rice defense responses, we used the CRISPR/Cas9 system to knockout (*ko*) the endogenous *OsNAC2* and *OsEREBP1* genes in the WT line (Nipponbare). We generated two *ko* mutants (*osnac2 oserebp1-1* and *osnac2 oserebp1-2*) in the T_1_ generation, whose target sites were missing one and two bases, respectively (Supplemental Fig. S7). Figure 1 illustrated the negative regulatory effect of OsNAC2 on immunity and OsNAC2-Cas9 lines also shows strong disease resistance (Fig. 6A). Then, we investigated the disease phenotype of the *osnac2 oserebp1* mutants. Compared with the WT line, the *osnac2 oserebp1* mutant lines were more susceptible to PXO71 (Fig. 6A), and lesion lengths of up to 13 cm after two-weeks inoculation (Fig. 6B). Furthermore, the accumulation of ROS in *osnac2 oserebp1* after DAB and NBT staining were the least. (Fig. 6C). In addition, the expression of resistance marker genes *OsPR1a* and *OsPR10a* were significantly reduced than the WT (Fig. 6, D and E).

**Figure 6.**
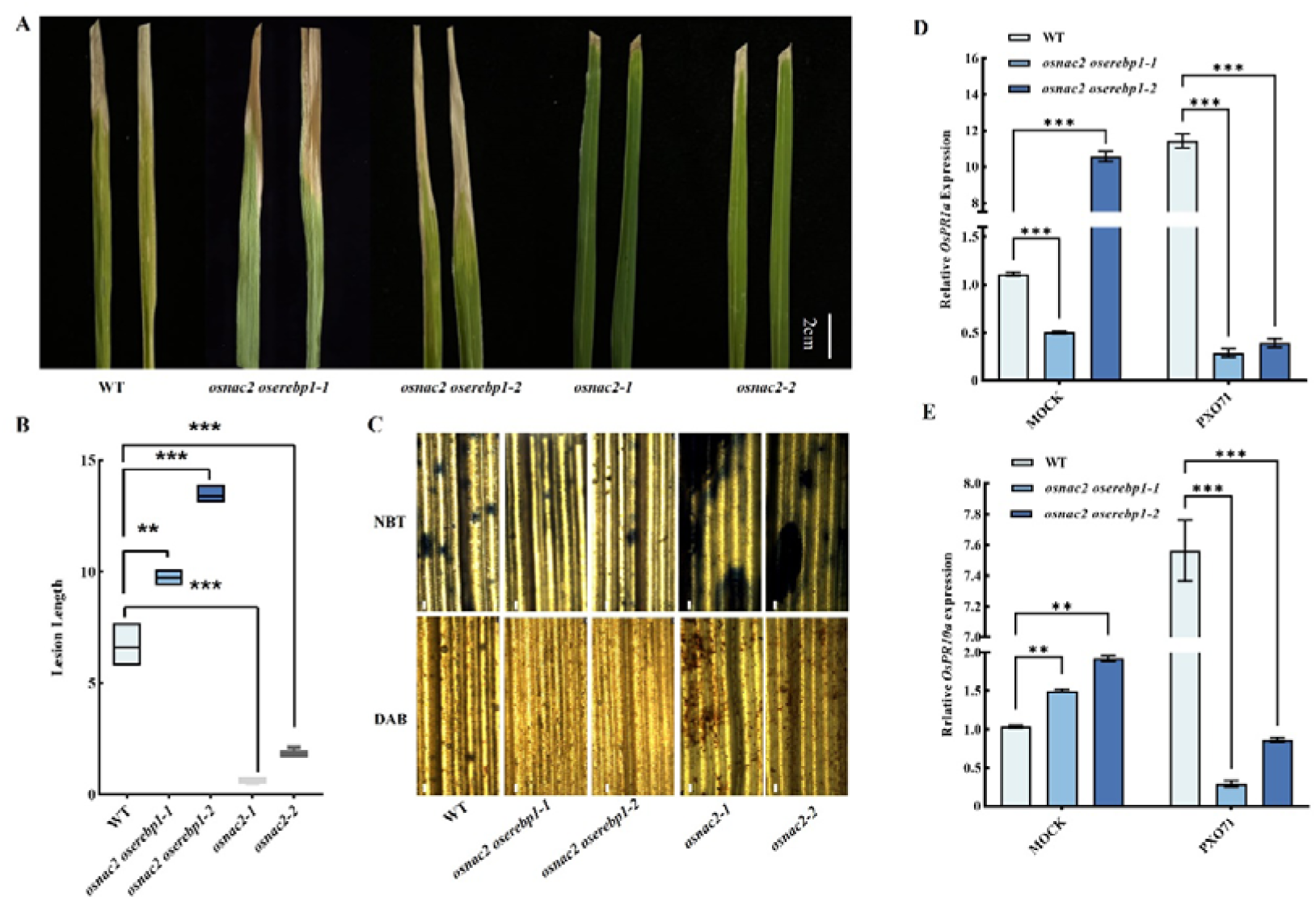
Verification and phenotype of *osnac2 oserebp1* mutant lines. A, Disease phenotype in the WT, *osnac2 oserebp1* mutant lines and OsNAC2-Cas9 lines after 4 weeks of infect with PXO71 under natural conditions. B, Lesion length. Scale bar=5 cm. D, NBT or DAB staining of the WT and *osnac2 oserebp1* mutant lines leaves after 4 weeks of infect with PXO71 under natural conditions. Scale bars=100 µm. E-F, Expressions of *OsPR1a* (E) and *OsPR10a* (F) genes in the WT and *osnac2 oserebp1* mutant lines under the *Xoo* strain PXO71 infection. Data are presented as the mean ± standard error of at least three biological replicates. Asterisks indicate significant differences between treatment and control by *t*-test. *P< 0.05, **P< 0.01, and ***P< 0.001.

That result indicated that *OsNAC2* negatively regulates disease resistance, while *OsEREBP1* positively regulates disease resistance through OsXb22a-mediated (Jisha et al., 2015). Therefore, both *OsNAC2* and *OsEREBP1* mediate rice resistance, but *OsEREBP1* may be more powerful than *OsNAC2*.

## Discussion

Effective defense responses must be induced in plants following pathogen infections (Lee and Dean, 1993). In response to a pathogen attack, plants perceive pathogen-derived signals and often activate complex defense signaling pathway networks (Peng et al., 2018). However, during long-term co-evolution, pathogenic bacteria developed the ability to suppress plant immune systems by modulating the complex hormone network (Xu et al., 2019). As an important plant defense hormone, SA plays a significant role in plant defenses, especially systemic acquired resistance (Vlot et al., 2009). After the pattern recognition receptors on the surface of plant cells recognize the PAMPs conserved among microorganisms, SA production is induced, leading to the establishment of PTI (Tsuda et al., 2010). Thus, the activation of SA production helps protect plants from pathogens during the early infection stage. Additionally, the key downstream regulator NPR1 enhances the responsiveness of disease-related genes (Zhang and Cai, 2005). In this study, the transcription factor OsNAC2 bound to the SA biosynthesis gene promoter, thereby affecting SA accumulation (Fig. 2), implying that OsNAC2 likely influences the PTI of plants.

Arabidopsis contains two *ICS* genes (*ICS1* and *ICS2*) as well as four *PAL* genes (*PAL1*–*PAL4*) (Huang et al., 2010), reflecting the importance of SA biosynthesis pathways for the survival of plants during evolution. The importance of the PAL pathway for SA biosynthesis under biotic and abiotic stress conditions remains to be determined (Zhang et al., 2021). Similarly, the combined effects of PAL and certain transcription factors on plant immune responses will need to be more thoroughly investigated (Kim and Hwang, 2014). Especially revealed that OsNAC2 may directly binds to the *OsICS1, OsPAL3 and OsPAL6* promoter in this study, and affect their transcription through two synthetic pathways, which inhibits the synthesis of SA and leads to the weakening of disease resistance in plants.

The coordinated nuclear and cytoplasmic activities of transcription factors are essential for the innate immune responses of plants to pathogens (García et al., 2010). How plants allocate specific proteins in the cytoplasm and nucleus following a pathogen infection to initiate appropriate defense responses will need to be elucidated. In this study, we observed that the nuclear distribution of OsEREBP1 is partially regulated by OsNAC2 through protein–protein interactions. OsEREBP1 has several signaling pathways in the nucleus to mediate immunity. For example, OsBWMK1, located in the nucleus, mediates the expression of *PR* genes by activating OsEREBP1 transcription factor and enhances resistance to pathogen infection (Koo et al., 2009).

Therefore, proteins that are not degraded by OsNAC2 in the nucleus can activate other signaling pathway. However, the silencing of *OsNAC2* decreased the accumulation of OsEREBP1 in the nucleus, the total abundance of OsEREBP1 increased, implying that the effects of OsEREBP1 on plant immunity are not restricted to its function in the nucleus. It is possible that a cytoplasmic ETI-related process promotes the accumulation of OsEREBP1. However, this possibility will need to be experimentally verified.

A previous study proved that the AP2/ERF multigene family can be divided into two subfamilies (AP2 and EREBP) according to the number of conserved domains (Chen et al., 2016). The EREBP subfamily members mainly participate in plant responses to organisms and environmental conditions, and most of them enhance plant disease resistance when overexpressed (Jisha et al., 2015). In the current study, we revealed that OsEREBP1 is functional in the nucleus and cytoplasm under normal conditions. Specifically, in the nucleus, OsEREBP1 inhibits the negative regulatory mechanism of OsNAC2 (Fig. 3) or it may combine with some GCC boxes (Iwamoto and Takano, 2011) and regulate transcription. In the cytoplasm, OsEREBP1 modulates stress resistance. For example, its interaction with OsXb22a influences bacterial blight resistance (Fig. 4). We demonstrated that the disease resistance of the *osnac2 oserebp1* double mutant was inferior to that of the WT control plants (Fig. 6), which confirmed that OsEREBP1 positively regulates the rice immune response. During the early stages of pathogen infection, OsNAC2 is highly abundant in the nucleus. The OsNAC2–OsEREBP1 interaction confines OsEREBP1 to the nucleus.

Some of the OsEREBP1 required for this interaction may be recruited from the cytoplasm. The large amounts of OsEREBP1 retained in the nucleus are degraded by the induced expression of *OsNAC2*, thereby limiting the effects of OsEREBP1 in the nucleus and enabling OsNAC2-mediated immune responses, including the inhibition of SA biosynthesis and the expression of disease resistance-related genes. When plants are continuously stimulated, the OsNAC2 content returns to its normal level and the OsEREBP1 function is no longer restricted. Consequently, plant defense activities are optimized.

On the basis of the study results, we established a model for the OsNAC2 function related to rice immunity (Fig. 7). Specifically, OsNAC2 directly regulates the expression of *OsICS1*, *OsPAL6* and *OsPAL3* to inhibit SA biosynthesis and signal transduction. Additionally, OsEREBP1 can interact with OsNAC2 in the nucleus and disrupts OsNAC2’s suppression on the *OsICS1*, *OsPAL3*-targeted genes expression, which balances the negative regulatory effects of OsNAC2 on rice disease resistance. However, *OsNAC2-*overexpression lines cause OsEREBP1 to accumulate in the nucleus. Following a pathogen infection, the OsEREBP1 confined to the nucleus is degraded because of the induced expression of *OsNAC2*, which interferes with the original homeostatic mechanism and increases the susceptibility of the rice to the pathogen.

**Figure 7.**
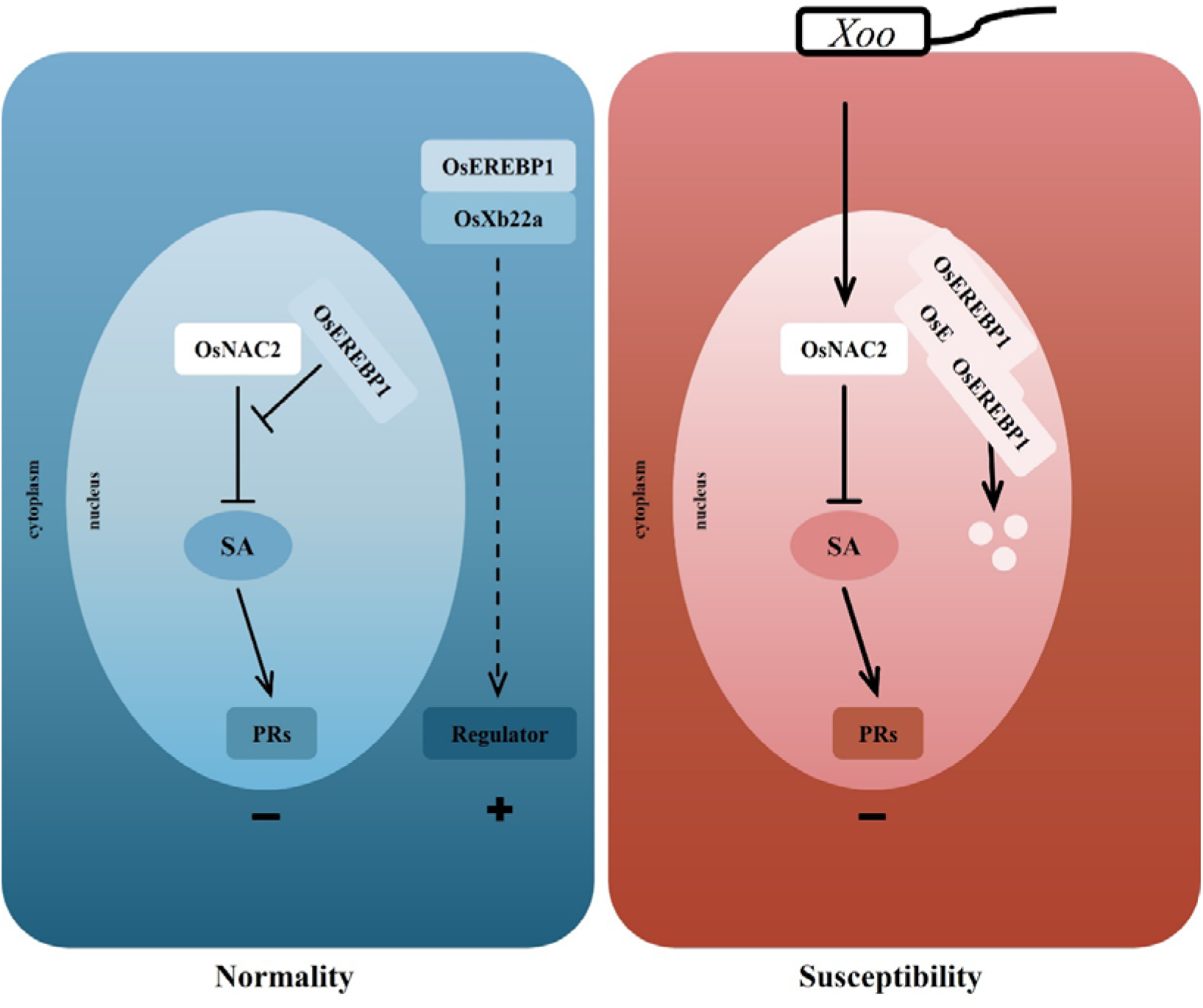
A model shows the OsNAC2 regulation module. Under normal circumstances, OsNAC2 and OsXb22a maintain a steady-state balance through OsEREBP1; when bacterial blight races infected, it activates the expression of OsNAC2, strengthens the inhibition of the SA signaling pathway and imprisons the OsEREBP1 protein in the nucleus to degradation, disrupting the homeostasis and causing plant susceptibility.

In summary, our research revealed the steady-state regulatory effects of OsNAC2-OsEREBP1 on the rice immune response. Enhancing the disease resistance of plants will lead to increased crop yields (Wang et al., 2021). We confirmed that the rice transcription factor gene *OsNAC2* may be useful for the genetic engineering of rice plants to generate high-yielding lines. Our findings also indicate that *OsNAC2*-RNAi plants are more disease resistant than WT plants (Fig. 1, B and C). Down-regulating *OsNAC2* expression can increase the grain yield by about 10% (Mao et al., 2017). The data presented herein may be of great significance for future studies aimed at improving rice disease resistance and crop yield through OsNAC2 production.

## Conclusions

In this study, we have confirmed that the transcription factor OsNAC2 and OsXb22a respectively compatibled with OsEREBP1 in the nucleus and cytoplasm, so as to realized the negative and positive regulation function of rice bacterial blight and exerted the immune homeostasis mechanism. When pathogenic bacteria infected, the induced expression of OsNAC2 leaded to OsEREBP1 confinement in the nucleus to degradation, resulted in the imbalance of homeostasis and the reduction of disease resistance in rice. So we established a model of the OsNAC2-OsEREBP1 homeostasis mechanism, which provided clues for the control of rice bacterial blight.

## Materials and methods

### Plant materials and treatments

Plant materials used in this study were previously generated *OsNAC2*-overexpressing lines (ON7 and ON11, expressing the *OsNAC2*-mGFP fusion protein) and *OsNAC2*-knockdown lines (RNAi25 and RNAi31) with Nipponbare genetic background (Chen et al., 2015; Mao et al., 2017). The rice seedlings were cultivated for 37℃ 16 hours during the day and 28℃ 8 hours at night in the incubator. In the experiment of inoculation of rice leaves, the tip part of mature rice leaves was cut off, and the exposed wound part was immersed in the bacterial solution (*Xanthomonas oryzae* pv. *oryzae*) for several seconds.

### RNA extraction and Real-time quantitative PCR (RT-qPCR) analysis

The total RNA was extracted from the (infected and uninfected) rice leaves with TsingZol Total RNA Extraction Reagent (Tsingke Biotechnology). The purified total RNA was reversed into the first-strand cDNA with Hifair III 1st Strand cDNA Synthesis SuperMix (Yeasen). The qPCR was performed using TSINGKE Master qPCR Mix (Tsingke Biotechnology) and the MyiQ2 Real-time PCR Detection system (Bio-Rad) according to the instructions of the manufacturer. The qPCR detection was repeated three times for each sample. All analyses used *OsActin* as the housekeeping gene. The primers used are listed Supplemental Table S1.

### NBT staining and DAB staining

The mature rice seedlings were treated with PXO71 for two weeks. The infected leaves were collected and stained with NBT (Coolaber) staining solution (50 mg NBT in 100 mL of 50 mM phosphate buffer pH7.8) or DAB (Tsingke Biotechnology) staining solution (1 mg mL^-1^ DAB) for 24 h at 28°C in the dark place. The leaves were decoloured in a boiling water bath in 90% ethanol and then preserved in 50% glycerol.

### Yeast two-hybrid (Y2H) assays

Because of OsNAC2 contains self-excitation domain, OsNAC2-N-terminal region coding region was amplified by PCR and subcloned into pGBKT7 prey plasmid. Coding regions of full-length and truncated OsEREBP1 were amplified and cloned into pGADT7 bait plasmid. Yeast strain Y2H Gold was co-transformed with specific bait and prey constructs through a lithium acetate-mediated method. The same method is used to determine the interaction relationship between OsEREBP1 and OsXb22a. Interactions were tested using SD/-Trp/-Leu/-His/-Ade medium. All primers for cloning these vectors were shown in Supplemental Table S2.

### Bimolecular fluorescence complementation (BiFC) assays

For the BiFC assay, four leaf stage *N. benthamiana* was selected as the experimental material. The coding regions of OsNAC2, OsEREBP1 and OsXb22a were amplified and cloned into nYFP and cYFP to generate OsNAC2-nYFP, OsEREBP1-cYFP and OsXb22a-nYFP, separately. Agrobacterium GV3101 was used to transform the recombinant vector. After activation, the bacteria were resuspended in the infection solution to infiltrate the *N. benthamianao* leaves (10 mM MES, 10 mM MgCl_2_ and 200 µM acetosyringone). Two days later, the injected leaf was observed using a confocal laser scanning microscope. The primers used are listed in Supplemental Table S3.

### Transient Transcription Dual-Luciferase Assay

For analyze the deactivation expression of *N. benthamianao* leaves, the sequence of 2.0 kb upstream from the ATG codon of the *OsICS1*, *OsPAL3 and OsPAL6* were cloned into pGreenII-LUC vector to generate the *LUC*/*REN* reporter genes, respectively. The OsNAC2 effector, OsEREBP1 effector or GFP control with reporter genes were co-transfected into *N. benthamianao* leaves, and the inoculated leaves were sampled after 48 hours. Use Dual Luciferase Reporter Gene Assay kit (Beyotime) to detect luciferase activity. The primers used are listed in Supplemental Table S4.

### Protein extraction and Western blot analysis

Total proteins were extracted using extraction buffer (10 mM NaCl, 0.25 mM MgCl_2,_ 1 mM Tris-HCl (pH 7.5), 0.25 mM DTT, 10% Tween, 0.5 tablet protease inhibitor). Protein samples were separated in a 10% SDS-PAGE gel and transferred to a PVDF membrane (66543, Pall Gelman Laboratory). Immunodetection of OsNAC2, OsEREBP1 and OsXb22a were performed using anti-HA, Myc or Flag primary antibody (1:5000) and secondary antibody anti-mouse IgG (1:5000). The corresponding molecular weight for OsNAC2, OsEREBP1 and OsXb22a in western blotting was consistent with the expected size. The anti-OsEREBP1 was from Beijing Huada Protein R & D Center Co., Ltd. The total protein and nucleoprotein were extracted from different transgenic strains of WT and NAC2 for antibody (1:5000) Western blot detection, and a single band size was detected at 33kDa.

### Co-immunoprecipitation (Co-IP) assays

In order to detect the interaction between OsNAC2 and OsEREBP1, OsXb22a and EREBP1, they were cloned on the PEG104 series of vectors to form PEG104-OsNAC2-HA, PEG104-OsEREBP1-Myc, PEG104-OsXb22a-Flag. The recombinant vector is on GV3101 transformation, transiently co-expressed in one-month-old *N. benthamiana* leaves, sampled after two days, extracted protein with extraction buffer, fixed on a rotator at 4℃ for 1h, filtered through 0.22 mm microporous expression PES membrane, take 400 µ as input, add the remaining sample to Myc magnetic beads and combine overnight, collect the magnetic beads on the magnetic stand, wash the magnetic beads three times with washing buffer: 10 mM NaCl, 0.25 mM MgCl_2,_ 1 mM Tris-HCl (pH 7.5), 0.25 mM DTT, 10% Trition-X, 0.5 tablet protease inhibitor, add protein loading and boil, and boil the input samples together, PEG104-Myc is the negative control. Use mouse monoclonal anti-HA, Myc or Flag antibody (1:5000) to detect the fusion protein. The primers are listed in Supplemental Table S5.

### Subcellular localization

For subcellular localization in rice protoplasts, cDNA fragment corresponding to the entire coding sequence of OsNAC2, OsEREBP1 and OsXb22a were cloned into the pRTV-GFP vector to generate pRTV-OsNAC2-cGFP (pUbi:OsNAC2-GFP), pRTV-OsEREBP1-cGFP (pUbi: OsEREBP1-GFP) and pRTV-OsXb22a-cGFP (pUbi: OsXb22a-GFP). The fusion construct was transformed or co-transformed with pSAT6:mCherry (RFP):VirD2NLS into protoplasts prepared from Nipponbare seedlings following the method as described previously (Bart et al., 2006). Fluorescence was examined under a confocal microscope (NiKon A1 i90, LSCM, Japan) at 16 h after transformation.

### Obtaining and Validation of *osnac2 oserebp1* double mutants

The knocking out mutants of *osnac2 oserebp1* bought from Baige Gene Technology Co., Ltd (Jiangsu, China). For mutation verification, we used two-week-old mutant and WT leaves of plants to extract DNA using the SDS method. The identification primers were provided by CAS to amplify the fragments of *OsNAC2* and *OsEREBP1* respectively (Supplemental Table S6). The obtained fragments were sequenced by Tsingke Biotechnology Co., Ltd. (Shanghai, China). In addition, TsingZol Total RNA Extraction Reagent (Tsingke Biotechnology) was used to extract total RNA from the leaves of the two-week-old lines to identify the expression levels of disease resistance-related genes.

### Liquid chromatography-tandem mass spectrometry (LC-MS/MS) analysis

The OsNAC2-GFP protein was expressed and purified as described in detail in GFP Co-IP assays. The purified OsNAC2-GFP recombinant protein was boiled for 10 min, separated by 12% SDS-PAGE. The protein was detected by coomassie brilliant blue (CBB) staining. The corresponding OsNAC2-GFP protein bands were sectioned according to the company’s requirements (APPLIED PROTEIN TECHNOLOGY). The protein was enriched and analyzed using Orbitrap Exploris™ 480 Mass Spectrometer (Thermo Scientific™). The mass spectrometry data were analyzed with Protein-Protein Interaction Networks (PPI) software.

### Statistical Analysis

All experiments were repeated at least three times in this study Data are presented as the mean ± standard error of at least three biological replicates, and the significant analysis was carried out with the Student’s *t*-test.

### Accession numbers

Sequence data from this article can be found in GenBank/EMBL databases under the following accession numbers: *OsNAC2* (Os04g0460600), *OsEREBP1* (Os02g0782700), *OsXb22a* (Os07g0171100), *OsActin* (Os10g0510000), *OsPR1a* (Os07g0129200), *OsPR10a* (Os12g0555500), *OsPAL1* (Os02g0626100), *OsPAL3* (Os02g0626600), *OsPAL6* (Os04g0518400), *OsICS1* (Os09g0361500), *OsNPR1* (Os01g0194300), *OsSGT1* (Os09g0518200) and *OsDSG1* (Os09g0434200).

## Acknowledgments

F. M. conceived the project and designed the study; Q. Z. and J. Y. performed the experiments; F. M. and X. M. provided technical assistance to Q. Z. and J. Y.; Q. Z. (major part) and J. Y. wrote the article; Q. Z. and F. M. revised the article; Y. W. modified all the figures. X. Y. provided mutant materials and related carriers for genetic transformation. All of the authors discussed the results and commented on the manuscript.

## Supplemental Data

The following supplemental materials are available.

Supplemental Figure S1. Phenotype of natural aging in OsRNAi31 and Nipponbare.

Supplemental Figure S2. Screening of interaction proteins of OsNAC2.

Supplemental Figure S3. Expressions of *OsSGT1*, *OsDSG1* and *OsEREBP1* in the WT and *OsNAC2*-transgenic lines.

Supplemental Figure S4. Analysis of NAC and ERF binding motifs in *OsICS1* and *OsPAL3* promoters.

Supplemental Figure S5. Western blot analysis of OsEREBP1 protein levels in cytoplasm and nucleus of the Nip.

Supplemental Figure S6. Identification of transgenic lines of *Ubi*: OsEREBP1-NLS and *Ubi*: OsEREBP1-NES.

Supplemental Figure S7. Mutation sequence of *osnac2 oserebp1* mutants.

**Supplemental Table S1.** Primers used for quantitative real-time PCR.

**Supplemental Table S2.** Primers used for yeast two hybrid.

**Supplemental Table S3.** Primers used for BiFC.

**Supplemental Table S4.** Primers used for LUC.

**Supplemental Table S5.** Primers used for Co-IP.

**Supplemental Table S6.** Primers used for mutant verification.

